# Perceptual salience is insufficient for auditory streaming in eastern gray treefrogs (*Hyla versicolor*)

**DOI:** 10.1101/2023.11.22.568351

**Authors:** Lata Kalra, Shoshana Altman, Mark A. Bee

## Abstract

Auditory streaming underlies a receiver’s ability to organize complex mixtures of auditory input into distinct perceptual “streams” that represent different sound sources in the environment. During auditory streaming, sounds produced by the same source are integrated through time into a single, coherent auditory stream that is perceptually segregated from other concurrent sounds. Based on human psychoacoustic studies, a prominent hypothesis regarding auditory streaming is that *any* perceptually salient acoustic difference between sounds can promote their segregation into distinct auditory streams. Here, we used the eastern gray treefrog, *Hyla versicolor*, to test this hypothesis in the context of vocal communication in a non-human animal. In this system, females choose their mate based on perceiving features of a male’s pulsatile advertisement calls in social environments (choruses) characterized by mixtures of overlapping vocalizations. We employed an experimental paradigm from human psychoacoustics to design interleaved pulsatile sequences (ABAB…) that mimicked key features of the species’ advertisement call, and in which alternating pulses differed in pulse rise time, which is a robust species recognition cue in eastern gray treefrogs. Using phonotaxis assays, we found no evidence that perceptually salient differences in pulse rise time promoted the segregation of interleaved pulse sequences into distinct auditory streams. These results suggest the hypothesis that any perceptually salient acoustic difference can be exploited as a cue for stream segregation is not supported in all species. We discuss these findings in the context of cues used for species recognition and auditory streaming.

## Introduction

Acoustic communication, and hearing more generally, frequently requires listeners to perceive relevant sound sequences as distinct from other concurrent sounds (Cherry & Taylor, 1954; McDermott, 2009). In humans, for example, following a conversation in noisy social settings (Remez, 2021; Repp, 1988) or recognizing a melody in an orchestral piece (Dowling, 2012; McDermott & Oxenham, 2008) involves the ability to hear sound sequences (e.g., words, syllables, musical notes) as distinct from other sounds occurring at the same time. The ability to hear distinct sound sequences amid competing sounds is a non-trivial challenge because sounds from multiple sources sum to form a composite sound wave that impinges on the ears of a listener (Bregman, 1990). The composite sound wave must be perceptually organized into distinct “streams,” each corresponding to a coherent representation of the sound sequence produced by a given source. This process, called “auditory streaming” (Bregman, 1990), involves two complementary processes in which sounds produced by the same source are *integrated* into a coherent auditory stream while sounds produced by different sources are *segregated* into separate streams (Bregman, 1990; Bregman & Campbell, 1971; Moore & Gockel, 2012).

Psychoacoustic studies in humans have uncovered various cues influencing the integration versus segregation of sounds during auditory streaming. Many of these studies have employed a simple experimental paradigm wherein subjects listen to interleaved sequences of two types of tone pulses (A and B) and report their perception of the rhythm or rate of the sequence. The acoustic differences between the A and B pulses are manipulated across trials. Integration versus segregation can be assessed using this ABAB stimulus paradigm to determine whether subjects report hearing, as a function of the acoustic differences between the A and B pulses, a single, integrated sequence (ABAB…) or two segregated sequences (A–A–… and B–B–…), each at half the pulse rate of the actual stimulus sequence (van-Noorden, 1974). Sufficiently large differences in the spectral content (e.g., fundamental frequency or timbre), temporal patterns (e.g., onset/offset times, amplitude and frequency modulation patterns) or spatial location of A and B sequences promote their segregation, while smaller differences are more likely to result in their integration (reviewed in Bregman, 1990; Darwin, 1997, 2008; Micheyl & Oxenham, 2010; Shamma et al., 2011). The breadth of acoustic cues that facilitate auditory streaming in humans led Moore & Gockel (2012) to hypothesize that *any* perceptually salient difference between sound sequences can promote their segregation into separate streams.

Many non-human animals communicate using rhythmic sequences of sounds, such as pulsatile calls in frogs and crickets (Prestwich, 1994; Gerhardt & Huber, 2002), song motifs in songbirds and whales (Winn et al., 1981; Bruno & Tchernichovski, 2019), and echolocation clicks in bats and dolphins (Nihoul, 2004). Moreover, these signals are perceived in complex acoustic environments consisting of multiple biotic and abiotic sound sources (Bee & Micheyl, 2008; Gerhardt & Huber, 2002; Greenfield, 2005). Auditory streaming is thus essential for accurate recognition, discrimination, and localization of signals across diverse species and behavioral contexts. Even though auditory streaming is a ubiquitous communication challenge, the phenomenon has so far received little attention in studies of non-human animal communication (Hulse, 2002; Bee & Micheyl, 2008; Dent & Bee, 2018). Preliminary investigations using the ABAB paradigm in non-human animals suggest similar auditory streaming cues are used in humans and a diversity of other species. Frequency differences, for example, promote segregation in insects (J. Schul & Sheridan, 2006), frogs (Nityananda & Bee, 2011), fish (Fay, 2000; Fay, 1998), birds (MacDougall-Shackleton et al., 1998; Itatani & Klump, 2014; Dent et al., 2016), and mammals (Izumi, 2002; Ma et al., 2010; Noda et al., 2013; Christison-Lagay & Cohen, 2014). Temporal differences in onset/offset times and amplitude modulation patterns promote segregation in frogs (Gupta & Bee, 2020) and birds (Itatani & Klump, 2009). Differences in spatial location promote segregation in insects (Dagmar von Helversen, 1984; Weber & Thorson, 1988), frogs (Farris et al., 2002; Farris et al., 2005; Bee, 2010) and mammals (Middlebrooks & Bremen, 2013; Yao et al., 2015). While these studies establish interesting parallels between auditory perception across taxa, it remains to be tested whether perceptual salience *per se* (sensu Moore & Gockel, 2012) is sufficient to promote segregation of sounds in non-human animals.

In this study of the eastern gray treefrog, *Hyla versicolor*, we used the ABAB stimulus paradigm to test the hypothesis that perceptually salient acoustic differences promote auditory streaming. The eastern gray treefrog is a well-studied frog in the context of animal communication that breeds in ponds and wetlands distributed throughout eastern North America (Gerhardt, 2001). Males of *H. versicolor* produce pulsatile advertisement calls (Fig. 1a) and breed in choruses. Even in small choruses of only conspecifics, there is a high degree of call overlap among neighboring males (Schwartz et al., 2002). In mixed-species choruses heterospecific males, including those of a morphologically indistinguishable sister species, *Hyla chrysoscelis*, also produce spectrally and temporally overlapping pulsatile advertisement calls (Fig. 1b) (Nityananda & Bee, 2011). Auditory streaming is thus crucial for female frogs to perceive the signal of a potential mate amidst other concurrent sounds (Bee, 2015). In *H. versicolor*, each advertisement call comprises of a sequence of 11-25 pulses (Fig. 1a). The amplitude time envelope of each pulse has a slow (approximately 65% of pulse duration) rise from pulse onset to peak amplitude and a fast (approximately 35% of pulse duration) fall from peak amplitude to pulse offset (Fig. 1c) (Gerhardt & Doherty, 1988; Ptacek et al., 1994; Gupta et al., 2021). Pulse amplitude rise and fall patterns – together described as “pulse shape” – facilitates species recognition in *H. versicolor*. Females of *H. versicolor* collected from a population in the State of Missouri in the central United States prefer pulses shaped with slow rise times, typical of conspecific calls (Fig. 1c), over pulses that have faster rise times and an overall shape that more closely resembles the heterospecific pulses of *H. chrysoscelis* (Fig. 1d). Rise time differences as small as 5 ms were perceptually salient and elicited strong behavioral discrimination between signals (Gerhardt & Schul, 1999). Here, we capitalized on this expected pulse rise time discrimination in *H. versicolor* to test the hypothesis that a perceptually salient difference in pulse rise time promotes the segregation of interleaved pulse sequences into separate auditory streams.

**Fig. 1.**
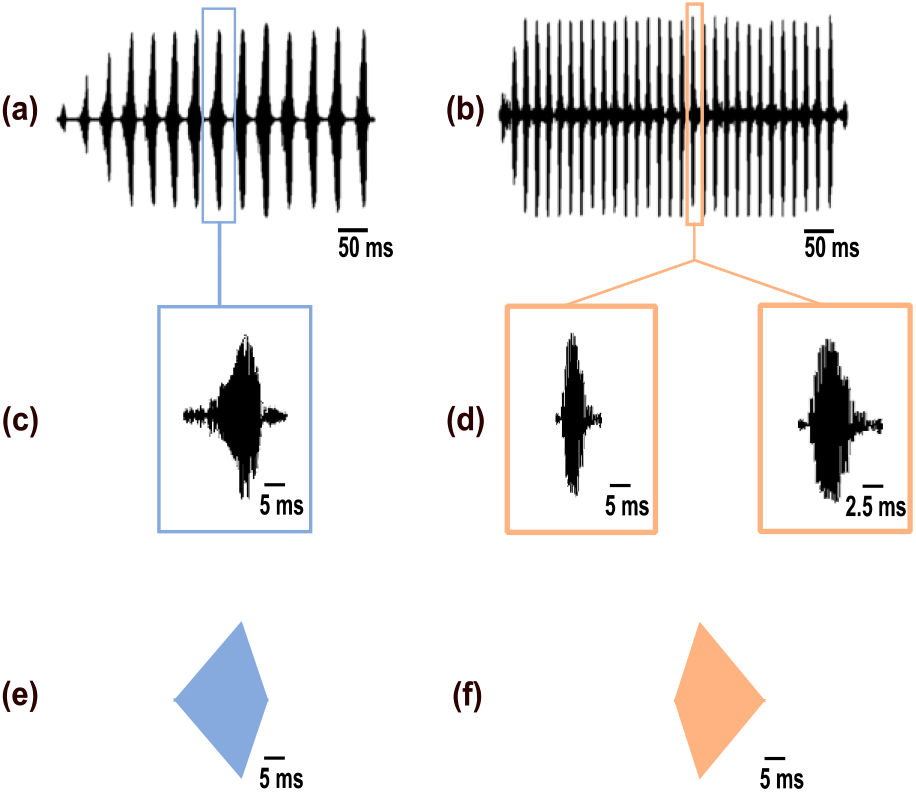
Natural and synthetic signals of *H. versicolor* and *H. chrysoscelis*. **a** Oscillogram of a natural advertisement call of *H. versicolor*. **b** Oscillogram of a natural advertisement call of *H. chrysoscelis* depicting a faster pulse rate compared to *H. versicolor*. **c** A highlighted natural pulse of *H. versicolor* depicting a slow rise and relatively faster fall in amplitude. **d** left: A highlighted natural pulse of *H. chrysoscelis* (shown in the same time-scale as *H. versicolor* in panel c) depicting a fast rise and relatively slow fall in amplitude, and right: The same pulse magnified two-fold to highlight how the pulse shape (relative rise and fall patterns) in *H. chrysoscelis* is almost reversed relative to that of a natural *H. versicolor* pulse. **e** Synthetic “A” pulse (in blue) modelled on the overall duration and the rise and fall-times of a natural *H. versicolor* pulse. **f** Synthetic “B” pulse (in orange), which is a digitally reversed version of “A” pulse, has an overall duration typical of a natural *H. versicolor* and an overall shape typical of a natural *H. chrysoscelis* pulse.

As a first step in our experimental design, we recorded and analyzed advertisement calls because pulse rise time preferences had not been established previously in our study population in the State of Minnesota. We used the results from our acoustic analyses to determine biologically realistic values of pulse rise time for subsequent behavioral experiments. We exploited a well-known preference for conspecific calls based on pulse-rate selectivity to design experiments based on the ABAB stimulus paradigm. The pulses in a *H. versicolor* call are, on average, about 20 ms long and separated by silent intervals of about 30 ms in duration. This regular rhythm corresponds to a pulse rate of 20 pulses/s (Gerhardt & Doherty, 1988; Gupta et al., 2021). Females of *H. versicolor* prefer the slower pulse rate of conspecific calls (Fig. 1a) over the faster pulse rate of *H. chrysoscelis* calls (Fig. 1b), which, on average, is approximately double (40 - 65 pulses/s) the pulse rate of conspecific calls (Noble & Hassler, 1936; Blair, 1958; Gerhardt, 1978; Ward et al., 2013). Accurate pulse-rate perception is crucial for species recognition as highlighted by the finding that two interleaved and identical conspecific pulse sequences are perceived as a single sequence with fast pulse rate that is less attractive to females of *H. versicolor* (Schwartz & Gerhardt, 1995; Schwartz & Marshall, 2006; see also Bee & Riemersma, 2008). Our use of the ABAB stimulus paradigm was based on a female’s pulse-rate selectivity. We broadcasted two interleaved sequences of pulses having the same conspecific pulse rate (each 20 pulses/s) but differing in their pulse rise time (A and B). The “A” pulses (Fig. 1e) had the pulse duration and rise time typical of conspecific pulses (Fig. 1c). The “B” pulses (Fig. 1f) were time-reversed versions of the “A” pulses, and therefore, had a pulse duration typical of conspecific pulses but an overall shape typical of heterospecific *H. chrysoscelis* pulses (Fig. 1d). We then measured stream segregation based on whether the subjects perceived two separate (A–A–… and B–B–…) sequences (indicating *segregation*), each with a preferred pulse rate (20 pulses/s), or a single (ABAB…) sequence (indicating *integration*) with a less preferred pulse rate (40 pulses/s) (Fig. 2). According to our hypothesis, we predicted subjects would be attracted to interleaved sequences that could be segregated into separate auditory streams, one of which was attractive, based on a perceptually salient difference in pulse rise time (Fig. 2).

**Fig. 2.**
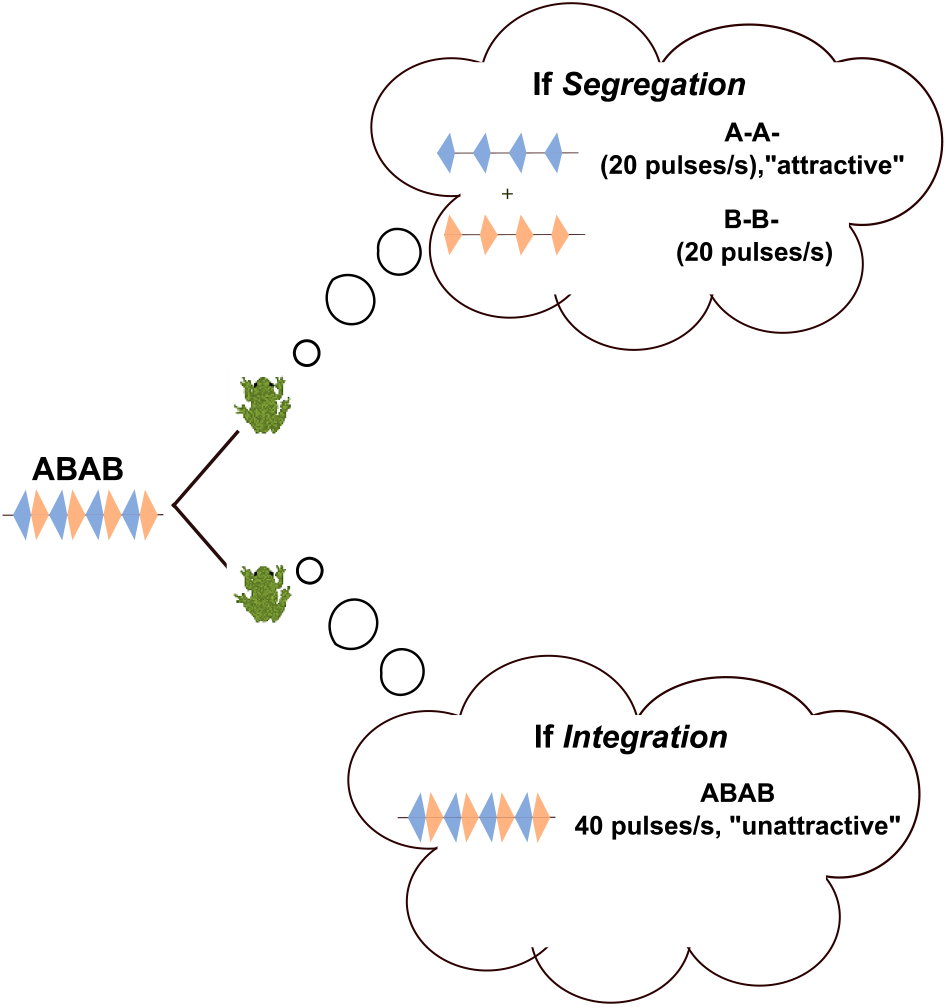
Protocol for testing auditory streaming. We broadcasted interleaved pulsatile sequences of A (in blue) and B (in orange) pulses (ABAB) to female *H. versicolor.* If pulse rise time differences were sufficient to promote segregation of sounds, we expected the females to perceive two distinct sequences, A–A– and B–B–, both of which had the preferred conspecific pulse rate of 20 pulses/s and one of which (A–A–) also had the preferred conspecific pulse rise time. Consequently, the ABAB stimulus was predicted to be attractive to females. In contrast, if the rise time differences between the A and B pulses were insufficient to promote segregation, we would expect females to perceive a composite ABAB sequence as having a relatively less attractive pulse rate of 40 pulses/s.

## Methods

### Subjects

All acoustic recordings and behavioral tests were conducted using subjects from the Tamarack Nature Center (Ramsey County, MN, USA), which belong to the Midwest clade of *H. versicolor* (Booker et al., 2022). Acoustic recordings of males (*n*=30) were made at night (between 2100 and 0100 h) in May and June of 2006 and 2021. For behavioral tests, females (*n*=43) were collected in amplexus at night (between 2100 and 0100 h) in May and June of 2021. Amplexed pairs were returned to the laboratory where they were maintained at approximately 4°C to delay egg laying and maintain behavioral responsiveness (Gerhardt, 1995). Prior to behavioral testing, frogs were placed in an incubator for at least 30 minutes and allowed to reach a body temperature of 20°C. Between trials, females were returned to the 20°C incubator with their mates for a minimum of 5 min to maintain body temperature and preserve responsiveness. Because *H. versicolor* breeds syntopically with *H. chrysoscelis* at our field site, we confirmed the species identity of all subjects in an initial two-alternative choice test in which we broadcasted alternating synthetic models of the two species’ calls (as in Gupta et al., 2021). Only females that approached the *H. versicolor* stimulus were used as subjects in the experiments described below. In some case, females were also used as subjects for other experiments not described here. There is little evidence for “carryover” effects between consecutive phonotaxis tests separated by several minutes (Akre & Ryan, 2010; Gerhardt, 1981). All frogs were released at their collection site within 48 hours of completing behavioral tests.

### Acoustic recordings and analysis

Vocalizations were recorded (44.1 kHz sampling rate, 16-bit resolution) using Sennheiser ME66 or ME67 microphones (Sennheiser USA, Old Lyme, CT, U.S.A.) connected to Marantz PMD620 or PMD670 recorders (D&M Professional, Itasca, IL, U.S.A.). Microphones were held by hand or mounted on a tripod, and the tip of the microphone was positioned approximately 1 m away from the focal male. For each individual male (*n* = 30) we recorded and analyzed a minimum of 5 calls (range, 5 - 45 calls/male). Since both the acoustic properties of advertisement calls and female preferences for call properties can vary with temperature (Gerhardt, 1978), we measured the wet-bulb air temperature and the water temperature at each male’s calling site immediately following each recording. We noted the general position from which the male was calling (e.g., in air on emergent vegetation versus floating on the surface of the water) to determine the most appropriate temperature for later use to standardize call properties to a common temperature of 20°C. We recorded males from two different ponds and from different areas within each pond across nights to reduce the chances of recording the same individual multiple times.

Acoustic recordings were analyzed using SoundRuler version 0.9.6.0 (Gridi-Papp, 2007), which performs automatic recognition of small repeated acoustic elements and exports an output summary of numerous acoustic properties (Bee, 2004). The output summary was further analyzed in R studio (R Core Team, 2020) to derive and analyze specific acoustic properties of interest for which we computed means, standard deviations (SD), and ranges. Our primary focus was on pulse shape, which we characterized by measuring pulse rise time (ms, time from pulse onset to peak amplitude) and pulse fall time (ms, time from pulse peak amplitude to offset). To place measures of pulse shape in the overall context of the advertisement call, we also measured other temporal properties, including pulse duration (ms), pulse rate (pulses/s), call duration (pulses/call), call rate (calls/min), and spectral properties including the frequency (Hz) of each pulse’s first and second harmonics, which correspond to the fundamental frequency and dominant frequency, respectively. Because the recordings were made at different temperatures, we followed Platz & Forester (1988) to standardize all call properties to 20°C, which is close to the average temperature observed in our recordings as well as the temperature at which we performed behavioral experiments.

### Acoustic stimuli

Synthetic acoustic stimuli (44.1 kHz sampling rate, 16-bit resolution) were generated in MATLAB R2020b (Mathworks, Natick, MA, USA) using parameter values taken from our acoustic analysis of natural signals. Across all experiments, stimuli were designed to stimulate a calling male and were constructed as a 5 min sequence of synthetic calls that repeated at a rate of 10 calls/min. Each call was generated as a sequence of pulses wherein each pulse was 20 ms long and composed of two phase-locked spectral components (1250 Hz and 2500 Hz, corresponding to the fundamental and dominant frequencies, respectively, of the natural signals). Further, based on our acoustic analysis, the amplitude of the 1250 Hz component was fixed to be 13 dB lower relative to the 2500 Hz component. The calls within each stimulus sequence differed in the rise time, rate, and timing of their constituent pulses according to the type of phonotaxis test performed, as described next.

### Experimental design

We performed three different choice tests, described below, using female phonotaxis as a behavioral assay. Each test was replicated twice using stimuli presented at one of two different sound pressure levels (80 dB peak SPL and 100 dB peak SPL). We used these sound pressure levels because auditory perception in frogs can be sound-level dependent (Gerhardt, 1987, 2005, 2008) and because these levels encompass much of the natural range of variation in the sound pressure levels of advertisement calls (Gerhardt, 1975). Each subject was tested in each of the six tests (3 choice tests × 2 signal levels) in a randomized order. We used two-tailed binomial tests to compare the proportion of females choosing a specific stimulus to the chance expectation if they chose randomly. All data analysis was performed in R studio (R Core Team, 2020).

### Perceptual salience test

We performed a two-alternative choice test to determine whether pulse rise time differences are perceptually salient in our study population. This test simulated a choice between two calling males producing calls having attractive pulse rates of 20 pulses/s, with each call comprising 16 pulses that differed only in pulse rise time (“A” versus “B”). “A” pulses (Fig. 1e) had slow, linear rise times (13 ms, 65% of pulse duration) and fast, linear fall times (7 ms, 35% of pulse duration). The “A–A–” stimulus (Fig. 3a) consisted of pulses that simulated the average rise time of conspecific pulses in *H. versicolor* as determined in our acoustic analysis (Noble & Hassler, 1936; Blair, 1958; Gerhardt, 1978). In contrast, the “B–B–” stimulus (Fig. 3b) consisted of time-reversed A pulses that had fast, linear rise times (7 ms, 35% of pulse duration) and slow, linear fall times (13 ms, 65% of pulse duration). These “B” pulses (Fig. 1f) closely resembled the overall shape of pulses in the heterospecific calls of *H. chrysoscelis* (fast pulse rise ∼ 35% of pulse duration and slow pulse fall ∼58% of pulse duration; Ward et al., 2013). By using time-reversed “A” pulses in the “B–B–” stimulus, we ensured both stimuli had pulses of consistent duration and peak sound pressure levels and differed only in pulse rise time and fall time (Diekamp & Gerhardt, 1995). Based on expectations from previous work by Gerhardt and Schul (1999) in Missouri populations, we predicted a proportion of subjects significantly higher than 0.5 would choose the A–A– stimulus (Fig. 4a). As discussed below, this prediction was supported by the data, thus allowing us to use the perceptually salient differences between the “A” and “B” pulses to test an auditory streaming hypothesis.

**Fig. 3.**
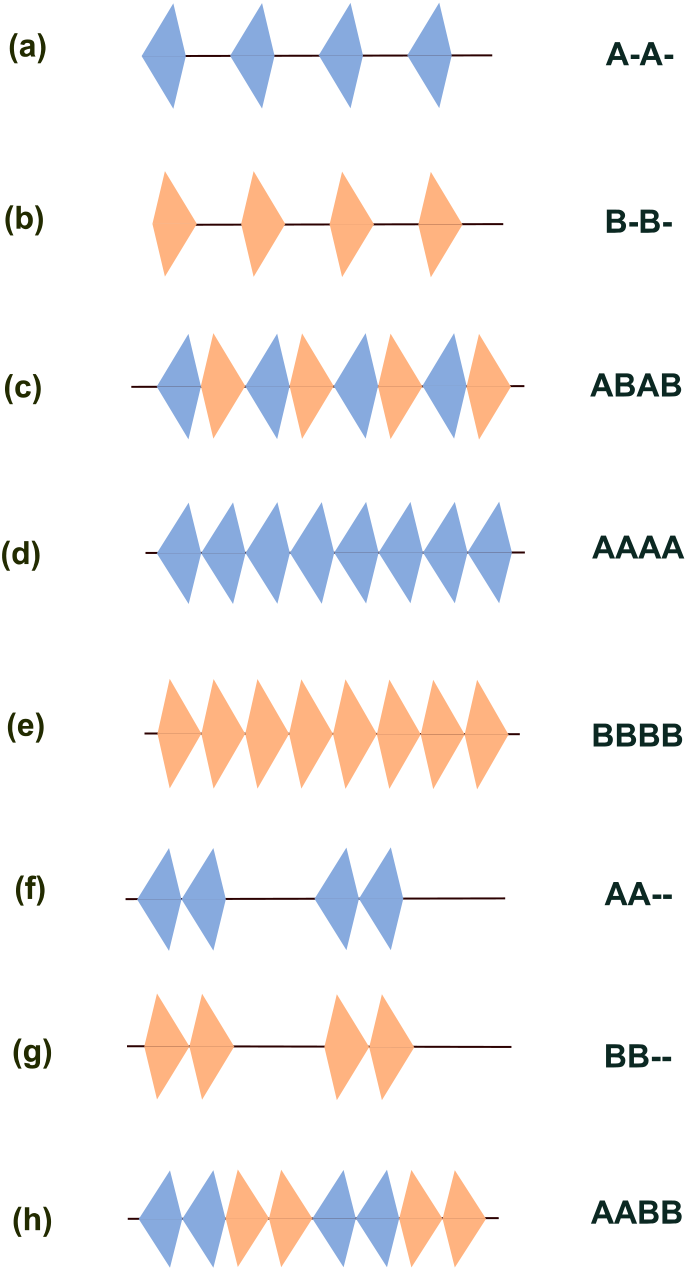
Acoustic stimuli for behavioral experiments. **a** “A-A-,” and **b** “B-B-” were constructed as sequences of A and B pulses, respectively, repeating at a rate of 20 pulses/s and having a regular inter-pulse interval of 30 ms. **c** “ABAB” was constructed by temporally interleaving the “A-A” and “B-B-” sequences and had a composite rate of 40 pulses/s. **d** “AAAA,” and **e** “BBBB” were constructed as sequences of A and B pulses, respectively, repeating at a rate of 40 pulses/s. **f** “AA—,” and **g** “BB—”, were constructed as sequences of A and B pulses, respectively, repeating at an average rate of 20 pulses/s and had an irregular inter-pulse interval that shifted between 10 ms and 50 ms between consecutive pulses. **f** “AABB” was constructed by temporally interleaving the “AA—” and “BB—” sequences and had a composite rate of 40 pulses/s.

**Fig. 4.**
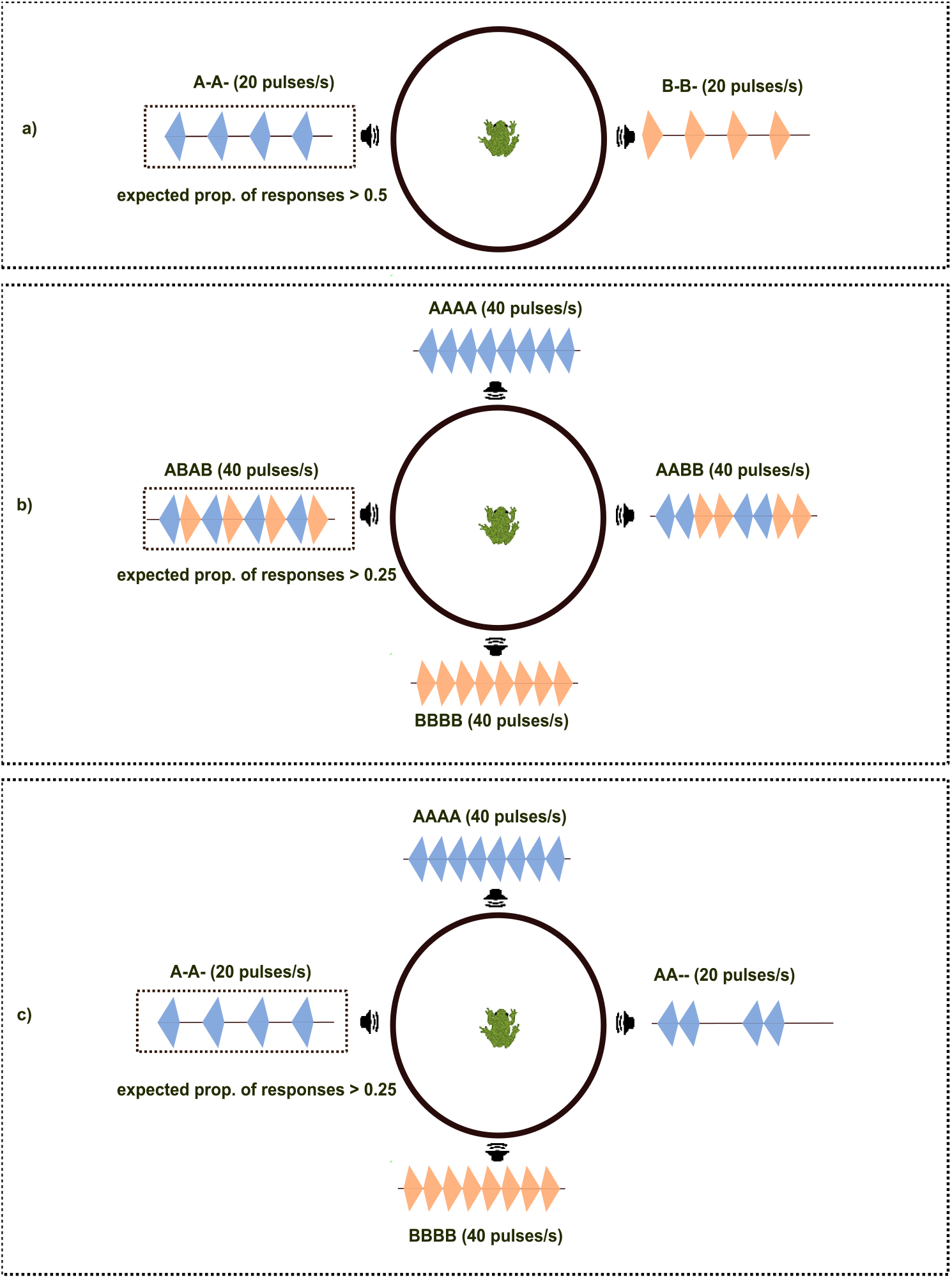
Design and predictions for the behavioral experiments. **a** Perceptual salience test. If the rise time differences between pulses A and B are perceptually salient, subjects were expected to prefer A-A-stimulus more than the chance probability of 0.5. **b** Auditory streaming test. If perceptually salient rise time differences are sufficient to allow auditory streaming, subjects were expected to prefer ABAB stimulus more than the chance probability of 0.25. **c** Pulse-rate and pulse-timing test. If subjects prefer calls with conspecific pulse rates and evenly spaced pulses, they were expected to prefer A-A-stimulus more than the chance probability of 0.25.

### Auditory streaming test

We used a four-alternative choice test to evaluate the hypothesis that females of *H. versicolor* can use perceptually salient difference in pulse rise time to segregate temporally overlapping calls into separate auditory streams. The key stimulus was based on the ABAB stimulus paradigm. It was created by temporally interleaving pulses from the A–A– and B–B– stimuli from the perceptual salience test to produce an “ABAB” stimulus (Fig. 3c). This ABAB stimulus had 32 pulses and a composite pulse rate of 40 pulses/s (simulating a less attractive *H. chrysoscelis* call) but was made up of two component pulse sequences (A–A– and B–B–) each having an attractive conspecific pulse rate of 20 pulses/s, only one of which (A–A–) also had the more attractive pulse rise time of conspecific calls. Two other stimuli in this four-alternative choice test also had the less preferred pulse rate of 40 pulses/s and consisted of a sequence of either all A pulses (“AAAA,” Fig. 3d) or all B pulses (“BBBB,” Fig. 3e). The final stimulus was created by interleaving pairs of A (“AA––,” Fig. 3f) and B (“BB––,” Fig. 3g) pulses so that, like the ABAB stimulus, this “AABB” (Fig. 3h) stimulus consisted of two pulse sequences having average pulse rates of 20 pulses/s. The main difference between the ABAB and AABB stimuli was that the former was comprised of two interleaved sequences having “regular” pulse timing (A–A– and B–B–), as determined by their constant 30 ms inter-pulse interval between consecutive pulses (typical of natural advertisement calls), whereas the component sequences in the AABB stimulus had “irregular” pulse timing (AA–– and BB––) created by having inter-pulse intervals that alternated between 10 ms and 50 ms between consecutive pulses but averaged to 30 ms over the duration of each composite stimulus. Among all the pulse sequences used across the four stimuli, the A–A– component sequence in the ABAB stimulus was expected to be the most attractive because it was the only stimulus with the pulse rise times, pulse rate, and pulse timing typical of conspecific calls (Gerhardt & Doherty, 1988; Gerhardt & Schul, 1999; Gerhardt, 2005). If the perceptually salient difference between the A and B pulse rise times was sufficient to allow auditory streaming, we predicted females would be attracted to the A–A– component of the ABAB stimulus and thus choose the ABAB stimulus over the other three stimuli, which had less preferred pulse rates (AAAA, BBBB, and AABB), pulse rise times (BBBB and AABB), or pulse timing (AABB). Therefore, if auditory streaming of the interleaved A and B pulses in the ABAB stimulus occurred, we predicted that the proportion of subjects choosing the ABAB stimulus would be significantly higher than 0.25 (Fig. 4b).

### Pulse-rate and pulse-timing test

We performed a final four-alternative choice test to confirm that females in our population were selective for conspecific pulse rates and regular pulse timing. The A–A– stimulus had a conspecific pulse rate of 20 pulses/s, the conspecific pulse rise time (A), and regular pulse timing. The AAAA and BBBB stimuli both had a faster pulse rate of 40 pulses/s (typical of the heterospecific calls of *H. chrysoscelis*) and regular pulse timing but differed in having either conspecific (A) or heterospecific (B) pulse rise times. Finally, the AA–– stimulus had a conspecific pulse rate of 20 pulses/s (on average), the conspecific pulse rise time (A), but irregular pulse timing (alternating 10 ms and 50 ms inter-pulse intervals). We predicted that if females prefer calls with conspecific pulse rates and evenly spaced pulses – two key provisions of our test of auditory streaming – then they would choose the A–A– stimulus at a rate significantly higher than the chance proportion of 0.25 (Fig. 4c).

### Testing protocol

Behavioral tests were performed in a 2-m diameter circular phonotaxis arena surrounded by a 60-cm tall wall. The arena wall was constructed from hardware cloth and black fabric to create a visually opaque but acoustically transparent barrier. The arena was set within a hemi-anechoic sound chamber (length × width × height: 2.8 × 2.3 × 2.1 m; Industrial Acoustics Company, IAC, North Aurora, IL, USA). Stimuli were broadcast from an HP ProBook 450 G6 (HP inc., Palo Alto, CA, USA) through a MOTU M4 sound card (MOTU, Inc., Cambridge, MA, USA) using Adobe Audition 3.0 (Adobe Systems Inc. San Jose, CA, USA). The output audio was amplified by a Crown XLS 1000 High-Density Power Amplifier (Crown International, Los Angeles, CA, USA) and played through one of four Orb1 speakers (Orb Audio, Sherman Oaks, CA, USA) located outside the arena wall on the floor of the sound chamber. The four speakers were evenly spaced and separated by 90° around the circumference of the circular test arena and positioned to face inward toward the center of the arena. The sound pressure level (SPL, LCpeak, re 20 µPa) of stimuli broadcast through each speaker was measured for calibration using a sound level meter (Larson Davis Model 831, Larson Davis Inc., Depew, NY) attached to a microphone placed at the center of the arena at the same level above the floor as a subject’s ears and aimed toward the speaker. For four-choice tests, alternative stimuli were broadcasted through four different speakers while for two-choice tests, alternative stimuli were broadcasted through two speakers located 180° apart. The consecutive calls within each stimulus sequence were separated by a silence of 5.23 s. The timing of calls across all stimulus sequences was such that an equal period of silence preceded and followed each call to avoid call overlap and any leader/follower relationships among stimulus calls. We further controlled for any leader/follower relationships by randomizing, across subjects, the order in which the very first calls of different stimuli were broadcast in a playback.

At the start of each test a single subject was placed at the center of the circular arena inside an acoustically transparent release cage. After a 60-s acclimation period, stimulus broadcasts began. After two repetitions of each test stimulus, a timer was started, and the frog was remotely released by lifting the lid of the release cage using a pulley system that could be operated from outside the sound chamber. Subjects’ responses were monitored through an overhead IR camera mounted directly over the test arena. Subjects were given up to 5 min to respond. A response was recorded if a subject approached to within 10 cm of a speaker and remained there for 30 s. A no-response was recorded if a frog failed to exit the release cage within 3 min after its release or if it failed to meet our response criterion within 5 min.

## Results

### Call analyses

The mean (± SD) rise and fall times of *H. versicolor* pulses were 13.0 ms (± 2.6 ms; range = 7.9 - 19.6 ms) and 7.4 ms (± 1.8 ms; range = 4.4 - 14.1 ms), respectively. The mean pulse duration was 20.4 ms (± 3.1 ms; range = 13.8 - 26.8 ms). Thus, on average, the pulse rise and fall times, respectively, were close to 65% and 35% of the call duration. Descriptive statistics for all other acoustic properties are reported in Table 1. Based on these results, we chose the rise and fall times of “A” pulses as 13 ms and 7 ms respectively. Since “B” pulses were digitally reversed versions of “A” pulses, the rise and fall times of “B” pulses were 7 ms and 13 ms, respectively.

**Table 1:** Descriptive statistics of acoustic properties of *H. versicolor* advertisement calls (*n* = 30 males) recorded in Minnesota and standardized to a temperature of 20° C. The range of temperatures at which males were recorded was 10.2 – 29.0° C.

| Acoustic property | mean ± SD | Range |
| --- | --- | --- |
| Pulse rise time (ms) | 13.0 ± 2.6 | 7.9 - 19.6 |
| Pulse fall time (ms) | 7.4 ± 1.8 | 4.4 - 14.1 |
| Pulse rate (pulses/s) | 19.3 ± 3.1 | 14.9 - 25.3 |
| Pulse duration (ms) | 20.4 ± 3.1 | 13.8 - 26.8 |
| Call duration (pulses/call) | 15.9 ± 3.3 | 11.1 – 25.1 |
| Call rate (calls/min) | 14.1 ± 4.0 | 5.1 – 22.2 |
| Pulse fundamental frequency (Hz) | 1232.5 ± 85.5 | 1084.7 – 1501.2 |
| Pulse dominant frequency (Hz) | 2465.1 ± 170.9 | 2169.5 – 3002.4 |

### Perceptual salience test

In the two-alternative choice test comprising the perceptual salience test, approximately, 98% (42 out of 43) and 91% (39 out of 43) of the females tested responded by making a choice at sound pressure levels of 100 dB and 80 dB, respectively. We predicted that females would prefer signals having a conspecific pulse rate and slow pulse rise time (A–A–) over an alternative having a conspecific pulse rate but a fast pulse rise time, and overall shape typical of heterospecific *H. chrysoscelis* pulse (B–B–). The data were consistent with this prediction. The proportion of subjects choosing A–A– stimulus over the B–B– stimulus was significantly higher than expected by chance (0.50) at both signal levels. At 100 dB, approximately 95% of subjects chose A–A– (*n* = 42, *p* < 0.001) and at 80 dB, approximately 92% of subjects chose A–A– (*n* = 39, *p* < 0.001) (Fig. 5a). The observed behavioral discrimination based on pulse rise time confirmed that differences in pulse rise time were both perceptually and behaviorally salient.

**Fig. 5.**
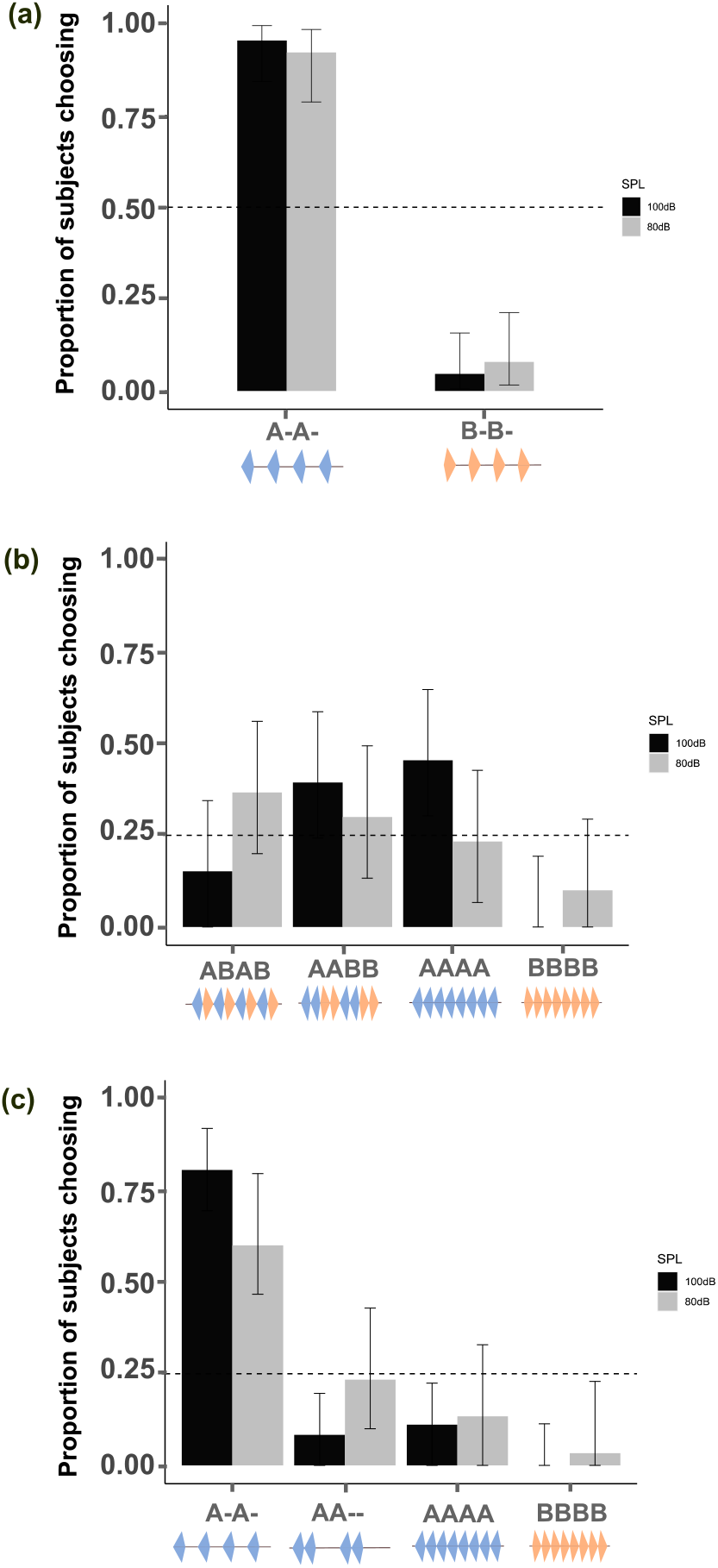
Results for behavioral experiments. Black and gray bars indicate the proportions of subjects choosing a given stimulus at 100 dB and 80 dB, respectively. **a** Results for the perceptual salience test. Error bars depict exact 95% binomial confidence intervals (CIs). **b** Results for the auditory streaming experiment. Error bars depict 95% multinomial CIs. **c** Results for the pulse-rate and pulse-timing test. Error bars depict 95% multinomial CIs. Horizontal dashed lines depict the chance probability for each experiment.

### Auditory streaming test

In the four-alternative choice test (ABAB vs. AAAA vs. BBBB vs. AABB) to investigate auditory streaming, approximately 77% (33 out of 43) and 70% (30 out of 43) of the females tested responded by making a choice at sound pressure levels of 100 dB and 80 dB, respectively. We predicted females would be attracted to the A–A– component of the ABAB stimulus if auditory streaming based on pulse rise time differences occurred. The data were not consistent with this prediction. The proportion of subjects choosing ABAB was not significantly higher than the chance probability of 0.25 (exact binomial test, α = 0.05) at either of the signal levels. At 100 dB, approximately 15% of subjects chose ABAB (*n* = 33, *p* = 0.231) and at 80 dB, approximately 37% of subjects chose ABAB (*n* = 30, *p* = 0.143) (Fig. 5b).

### Pulse-rate and pulse-timing test

In the four-alternative choice test (A–A– vs. AAAA vs. BBBB vs. AA––) to confirm pulse-rate and timing preferences, approximately 84% (36 out of 43) and 72% (31 out of 43) of the females tested responded by making a choice at sound pressure levels of 100 dB and 80 dB, respectively. We predicted that females would prefer the stimulus with a conspecific pulse rate and regular pulse timing (A–A–) over those with heterospecific pulse rates (AAAA and BBBB) and irregular pulse timing (AA––). The data were consistent with this prediction. The proportion of subjects choosing A–A– (slow rate and regular timing) was significantly higher than a chance probability of 0.25 (exact binomial test, α = 0.05) at both signal levels. The percentage of subjects choosing A–A– was approximately 81% at 100 dB (*n* = 36, *p* < 0.001) and 61% at 80 dB (*n* = 31, *p* < 0.001) (Fig. 5c).

## Discussion

The goal of this study was to test the hypothesis (sensu Moore & Gockel, 2012) that perceptual salience *per se* is sufficient to promote auditory streaming in non-human animals. Our results are inconsistent with this hypothesis. A species-typical difference in pulse rise time was perceptually salient, as evidenced by strong behavioral discrimination based on this acoustic cue in two-choice tests. However, there was no evidence that this salient acoustic cue also promoted the perceptual segregation of two interleaved pulses sequences differing in pulse rise time. Based on this outcome, we provisionally conclude that perceptual salience was insufficient for auditory streaming in the context of segregating temporal sequences of pulses in overlapping calls in *H. versicolor*.

Our bioacoustic analyses confirmed the presence of species differences in pulse rise time between *H. versicolor* and its sister species, *H. chrysoscelis*, in Minnesota that were similar to differences reported in other populations (Gerhardt & Doherty, 1988). Pulses in *H. versicolor* had rise times (range: 7.9 to 19.6 ms; Table 1) that were, on average, about 10 ms slower than those in the calls of *H. chrysoscelis* recorded in the same geographic area (range:1.8 to 4.7 ms; Ward et al., 2013). Moreover, our two-choice test of perceptual salience demonstrated that a rise time difference of just 6 ms was perceptually salient and elicited a robust preference (by 92 – 95% subjects) for slow rise times. This finding corroborates previous work on pulse rise time preferences in female *H. versicolor* from a Missouri population (Gerhardt & Doherty, 1988; Gerhardt & Schul, 1999). Both the absolute rise times between the A and B pulses used in our study (13 ms versus 7 ms, respectively), and their relative difference (6 ms) were close to those tested by Gerhardt & Schul (1999; e.g., 12.5 ms versus 7.5 ms). Our study used 20-ms pulses with two spectral components (1200 Hz and 2400 Hz), whereas Gerhardt & Schul (1999) used 25-ms pulses having just the lower or the higher spectral component alone. In both studies, females of *H. versicolor* rejected a fast rise time more typical of the pulses in calls produced by male *H. chrysoscelis* in favor of a slow rise time typical of the pulses in conspecific calls. Our findings add to the evidence that pulse rise time, along with other fine temporal features like pulse rate, facilitate pre-mating species isolation between *H. versicolor* and *H. chrysoscelis,* which have spectrally similar calls (Gerhardt, 2005). As such, the present findings also contribute to our current understanding of how signal preferences may persist or vary across different populations and geographical lineages of treefrogs (e.g., Gerhardt et al., 2007; Gupta & Bee, 2023).

Despite strong behavioral discrimination between two pulse sequences differing in pulse rise time (i.e. A–A– versus B–B–), there was no evidence that the same difference promoted auditory streaming when the same sequences were temporally interleaved (i.e., ABAB). Females did not prefer the ABAB stimulus when it was presented in a four-choice test with alternatives having less preferred pulse rates, pulse rise times, or pulse timing (AAAA, BBBB, and AABB). This result is contrary to our prediction that the rise time difference would promote segregation of the ABAB sequence into separate streams, one of which corresponded to a pulse sequence (A–A–) having the preferred pulse rate, pulse rise time, and pulse timing typical of conspecific calls. One alternative explanation for this lack of preference could be that subjects perceptually segregated the ABAB stimulus into separate A–A– and B–B– streams based on rise time differences but behaviorally avoided the source of the perceived B–B– stream consisting of pulses with fast rise times. This explanation seems unlikely for several reasons based on other work in this species. First, females of *H. versicolor* will approach the calls of a male *H. chrysoscelis* in a no-choice test when it is the only stimulus presented, suggesting stimuli with both fast pulse rates and fast pulse rise times are not inherently aversive (Gerhardt & Doherty, 1988). Second, Bush et al. (2002) and Schul & Bush (2002) showed that females responded in no-choice tests to a broad range of stimuli having different pulse rise times, including rise times faster than those of the B pulses in our stimuli. Third, Gerhardt et al. (1994) showed that females of *H. versicolor* did not avoid *H. chrysoscelis* calls while approaching a conspecific call, and Schwartz et al. (2000) showed that females of *H. versicolor* did not preferentially choose a conspecific call by itself over an identical alternative call that was paired with the call of a predator. Consistent with these findings, most females (≥ 70%) chose one of the four stimuli in our auditory streaming test (including the ABAB and AABB stimuli), which suggests B pulses were not inherently aversive. Results from an additional four-alternative choice test (see Supplementary Information) indicated B pulses can even be attractive in some stimulus contexts. Therefore, it seems highly unlikely that a perceived B–B– stream in the ABAB stimulus was in any way aversive in our test of auditory streaming. Finally, Stratman et al. (2021) demonstrated that females of *H. versicolor* preferentially approach small clusters of calling males over males calling in isolation. Had females perceptually segregated the ABAB stimuli into separate streams, one preferred (A–A–) and one less preferred (B–B–), then we might have expected the perceived presence of two males in close proximity to impart greater behavioral salience to the ABAB stimulus. Based on this other work, we interpret the lack of a significant preference for ABAB in our experiment as indicating that the pulse rise time differences did not promote auditory steaming.

Our study is the first investigation of the effects of pulse rise time differences on auditory streaming in a non-human animal. As such, our findings contribute to the existing knowledge on the effect of temporal differences on auditory streaming. While our study shows no effects of pulse rise time differences on auditory streaming, it would be worth testing the same hypothesis in other species, such as in some grasshoppers, which also use rise time as a behaviorally salient signal trait (Helversen, 1993). Besides our study, the only other investigations of the effect of amplitude rise time alone on segregation of sounds have been in humans. Similar to our study, Hartmann & Johnson (1991) tested the segregation of *sequential* sound elements and found rise time differences to be a weak facilitator of stream segregation. In that study, segregation of short (4 s) interleaved sequences of melodies (A and B) having different rise times was not any better than when melodies A and B had the same rise times. In contrast to our findings and those of Hartmann & Johnson (1991), Bregman et al. (1994a, b) demonstrated that rise time differences can facilitate segregation of sounds that occur simultaneously (as opposed to sequentially). In the studies by Bregman et al. (1994a, b), the discriminability of target tones in a multi-tone complex was better when the target exhibited a sudden rise compared to the other tones in the complex. Bregman et al. (1994a, b) speculated that a sudden onset or change in amplitude of target tones may “reset” the pitch-analysis mechanisms, leading to the segregation of target tones from the complex.

The apparent inability of rise time differences to promote sequential stream segregation in our study and that by Hartmann & Johnson (1991) must be considered in light of a well-known phenomenon in auditory streaming known as the “build up” effect. During segregation of *sequential* sounds, the percept of two distinct streams does not arise instantaneously but instead builds up over several seconds after stimulus onset (Bregman, 1978; Anstis & Saida, 1985; Micheyl et al., 2005; Deike et al., 2012). Behavioral measurements in humans (Anstis & Saida, 1985; Bregman, 1978; Thompson et al., 2011), ferrets (Ma et al., 2010) and budgerigars (Cai et al., 2018) demonstrate that when hearing interleaved tone sequences that differ acoustically, subjects initially perceive a single stream. The probability of perceiving two streams increases as the sequence progresses. This build-up of a two-stream percept over time has been attributed to the long-term adaptation of neural responses, as demonstrated in mammals (Micheyl et al., 2005; Snyder et al., 2006; Pressnitzer et al., 2008) and songbirds (Bee et al., 2010). Importantly, previous studies on the build-up of auditory streaming used long interleaved sequences (> 10 s) and found that the build-up of a two-stream percept took several seconds (5-10 s). In contrast, the study by Hartmann & Johnson (1991), which failed to find strong evidence for sequential stream segregation based on differences in rise time, used an overall stimulus duration that was relatively short at 4 s. While our study involved similar ABAB interleaved sound sequences, our stimulus design was constrained by the requirement to stimulate natural communication signals. Consequently, one limitation of our study is that it only examined auditory streaming over relatively short sequences of pulses within calls that were < 1 s in duration. It is primarily for this reason that our main conclusion, namely that salient pulse rise time differences do not promote stream segregation in gray treefrogs, must remain provisional. The ability of perceptually salient differences in pulse rise time to impact auditory streaming using longer stimulus sequences remains to be investigated in frogs.

Previous investigations of perceptual organization in treefrogs illustrate the importance of considering both stimulus design and the perceptual task. For example, previous studies of *H. chrysoscelis* using short, call-like sequences of pulses similar to those used in the present study have revealed the importance of common onsets/offsets (Gupta & Bee, 2020) and common spatial location (Bee, 2010) in promoting simultaneous integration of the two harmonics in the pulses of gray treefrogs calls. In contrast to the study by Bee (2010), the effect of spatial separation between consecutive pulses in short, call-like pulse sequences had markedly less impact on promoting sequential segregation in both *H. versicolor* (Schwartz & Gerhardt, 1995 and Schwartz & Del Monte, 2019) and *H. chrysoscelis* (Bee & Riemersma, 2008). This discrepancy in the strength of spatial separation as a segregation cue across sequential versus simultaneous segregation tasks parallels the contrast between findings on pulse rise time from the present study of frogs and those of humans by Bregman et al. (1994 a, b). One study of sequential segregation in *H. chrysoscelis* found that females could segregate a short, call-like sequence of pulses (A–A–) that was periodically interleaved with the pulses in a long (5 min) and continuous sequence of pulses (B–B–) differing in frequency, provided there was sufficient frequency separation between the A and B pulses (Nityananda & Bee, 2011). Whether pulse rise time differences might promote auditory streaming using a similar stimulus paradigm remains to be investigated.

Finally, it is also worth consider the lack of an effect of pulse rise time in the light of complex cue interactions during auditory streaming. In natural auditory scenes, multiple cues, or acoustic differences, are available to a receiver and may be differentially weighed during auditory streaming. For instance, Elhilali et al. (2009) tested auditory streaming in a cue conflict scenario using two sequences (A–A– and B–B–) that exhibited fairly large frequency separation, which promotes segregation, but shared coherent temporal onsets and offsets, which promotes integration. They found that coherent temporal onsets/offsets override frequency separation during auditory streaming as human subjects reported hearing a single stream (indicating integration). In other cases, different cues can also impact auditory streaming in an additive fashion. Micheyl et al. (2013), for example, found that inharmonicity (sounds having different fundamental frequencies) and temporal incoherence additively facilitate the segregation of sounds in humans. Importantly, while Hartmann & Johnson (1991) showed a weak effect of rise times on sound segregation (in the absence of frequency differences), Singh & Bregman (1997) showed an additive effect of rise times and frequency differences on stream segregation in humans. Non-human animals also incorporate cue interactions during auditory streaming, as seen for European starlings, Budgerigars and Zebra finches (Dent et al., 2016; Itatani & Klump, 2020). Additionally, there is also evidence for no interaction, as shown by Schwartz & Del Monte (2019) for spectral and spatial cues in *Hyla versicolor*. In the present study, A and B pulses had the same carrier frequencies, but different pulse rise times. It might be the case that spectral similarity overrides pulse rise time differences during auditory streaming. In such a case, we would expect that spectral similarity between A and B pulses promotes their integration irrespective of the differences in pulse rise times, which is in line with the findings of this study. Further, it might also be the case that differences in pulse rise time additively interacts with spectral differences to promote stream segregation. Additional studies that manipulate pulse rise time along with other potential cues, such as spectral differences or differences in spatial location, will be needed to uncover any interaction effects.

## Acknowledgements

We thank Saumya Gupta, Katie Krueger, Sophie Barno, Kaitlyn Bonnema, Sarah Bonnema, Olivia Groth, Rishi Gulati, Collin Meyer and Annika Ruppert for their help collecting and testing frogs, and Mike Goodnature from Ramsey County Parks and Recreation for generous access to collection sites. We are also grateful to Marlene Zuk, Andrew Oxenham, Katie Krueger and Satyabhama de Oliveira for feedback on this manuscript.

## Author contributions

LK and MAB: designed the study. LK and SA: conducted experiments. LK and SA: analyzed data. LK, SA and MAB: prepared the manuscript.

## Declarations

### Ethical approval

This research was approved by the University of Minnesota Institutional Animal Care and Use Committee (#0602A-81890 and #2001-37746A).

