## Supplemental Figure 1 for "Perceptual salience is insufficient for auditory streaming in eastern gray treefrogs (*Hyla versicolor*)"

### Methods:

Collection of subjects and the testing protocol is as decribed in the main text.

### Experimental design:

This four-alternative choice test was done to test the choice of females when encountering a trade-off between two key signal recognition features- pulse rise time and pulse rate. The two key alternative stimuli used here were, a 40 pulses/s sequence of A pulses (AAAA), which had an attractive pulse rise time but a less preffered pulse rate, and a 20 pulses/s sequence of B pulses (B–B–), which had an attractive pulse rate but a less preferred rise time. Besides the two key stimuli, we also used two other alternative stimuli- one, a 20 pulses/s sequence of B pulses (BB—), with both a less preferred rise time and pulse timing, and the other, a 40 pulses/s sequence of B pulses (BBBB), with both a less preferred rise time and pulse rate. We expected subjects to significantly prefer AAAA if pulse rise time overrides

### Results:

At 100 dB, females significantly preferred the AAAA stimulus (n= 38, *p*= 0.001) and at 80 dB females significantly preferred B–B– stimulus (n= 33, p < 0.001) (Fig. S1).

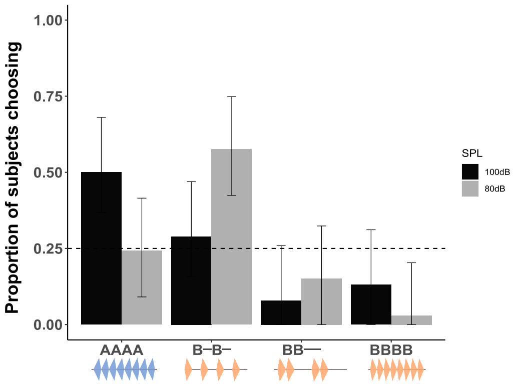


**Fig. S1** Results for the four-alternative choice test. Black and gray bars indicate the proportions of subjects choosing a given stimulus at 100 dB and 80 dB, respectively. Error bars depict 95% multinomial CIs. Horizontal dashed lines depict the chance probability for each experiment.
